# Tumor B cell infiltration in platinum-treated advanced urothelial carcinoma

**DOI:** 10.1101/2025.02.27.640395

**Authors:** Konrad Stawiski, Júlia Perera-Bel, Alejo Rodriguez-Vida, Núria Juanpere Rodero, Jihyun Lee, Daniel E. Michaud, Jennifer L. Guerriero, Kent W Mouw, Aristotelis Bamias, Filipe LF Carvalho, Joaquim Bellmunt

## Abstract

Platinum-based chemotherapy combined with immunotherapy provides durable disease control in advanced urothelial cancer. However, cisplatin and carboplatin differently impact the tumor immune microenvironment, affecting chemo-immunotherapy response. Here, we evaluate immune cell populations and ecosystems associated with overall survival in patients treated with platinum-based chemotherapy. Our transcriptomic analysis of pretreatment tumor samples from three cohorts (189 patients) of advanced urothelial cancer showed that lymphoid cell infiltration was significantly associated with prolonged overall survival. In cisplatin-treated patients, high memory B cell infiltration provided a significant overall survival improvement, but no such association was found in carboplatin-treated patients. Additionally, gene expression signatures implicated in B cell memory lineage and associated cytokines were associated with better overall survival in independent cancer patient cohorts. Our findings highlight memory B cell infiltration as a potential prognostic biomarker in urothelial cancer and emphasize the role of the tumor immune microenvironment in chemotherapy response.

## Introduction

Platinum-based chemotherapy followed by radical cystectomy has been the standard-of-care for muscle-invasive bladder cancer (MIBC) for the past three decades^1^. Randomized clinical trials^2^ and meta-analyses^3^ have shown that cisplatin-based neoadjuvant chemotherapy regimens are associated with improved overall survival (OS) compared with surgery alone. Interestingly, recent evidence from the NIAGARA Phase III clinical trial combining perioperative chemotherapy with immunotherapy demonstrated improved OS compared to chemotherapy alone.^4^ These results implicate the bladder tumor microenvironment (TME) as a key feature in response to perioperative systemic therapy and underscore the importance of understanding the interaction between chemotherapy and immunotherapy in bladder cancer.

A recent secondary analysis of the IMvigor130 clinical trial where patients with locally advanced and metastatic urothelial carcinoma were treated with either cisplatin or carboplatin in combination atezolizumab found that pre-existing adaptive immunity (evidence of an existing T cell-mediated immune response) in the TME was associated with improved response to immune checkpoint blockade^5^. Moreover, this analysis demonstrated that cisplatin, but not carboplatin, can modulate the tumor immune microenvironment and increase the efficacy of immune checkpoint inhibitors in patients with metastatic urothelial cancer ^5^. The interplay between cisplatin or carboplatin with the immune cell infiltrate as well as the impact of TME modulation on OS in urothelial cancer remain largely unexplored. Despite first-line platinum combination therapy being used less due to new data from phase III first-line studies, such the combination of enforumab vedotin (EV) and pembrolizumab^6^ the therapeutic sequence of cisplatin-based chemotherapy followed by switch maintenance immunotherapy is still considered and optional standard of care in areas where this EV plus pembrolizumab is not available.^7,8^ In this study, we analyzed transcriptional profiles of pre-treatment tumors from patients with locally advanced, surgically incurable, or metastatic bladder cancer who were treated with either cisplatin- or carboplatin-based chemotherapy alone. We assessed the association between OS and immune cell populations present in the primary tumor across three independent cohorts – a prospective randomized phase III trial (ACTRN 12610000845033), a real-world cohort, and a molecularly characterized MIBC cohort – to inform novel prognostic molecular signatures for patients with bladder cancer.

## Results

### Patient clinical characteristics

Patient and tumor characteristics are summarized in Table 1. Study group included 2 proprietary cohorts - the HM cohort (n = 55) and the GREEK cohort (n = 62), derived from a sub analysis of a phase III randomized clinical trial that excluded surgically eligible patients. Notably, the HM cohort included older and more medically fragile patients than the GREEK cohort (Table 1). Across both cohorts, the most frequently administered treatment regimen was a combination of cisplatin or carboplatin with gemcitabine (n = 80). In the HM cohort, 22 patients received carboplatin after being considered cisplatin-ineligible. Additionally, we used patients with locally advanced and metastatic cancer from the external TABER cohort (n = 72). The TABER cohort is a clinicogenomic publicly available cohort of bladder cancer patients from Taber et al. (Figure 1A). Our final study group comprised 189 patients.

**Figure 1.**
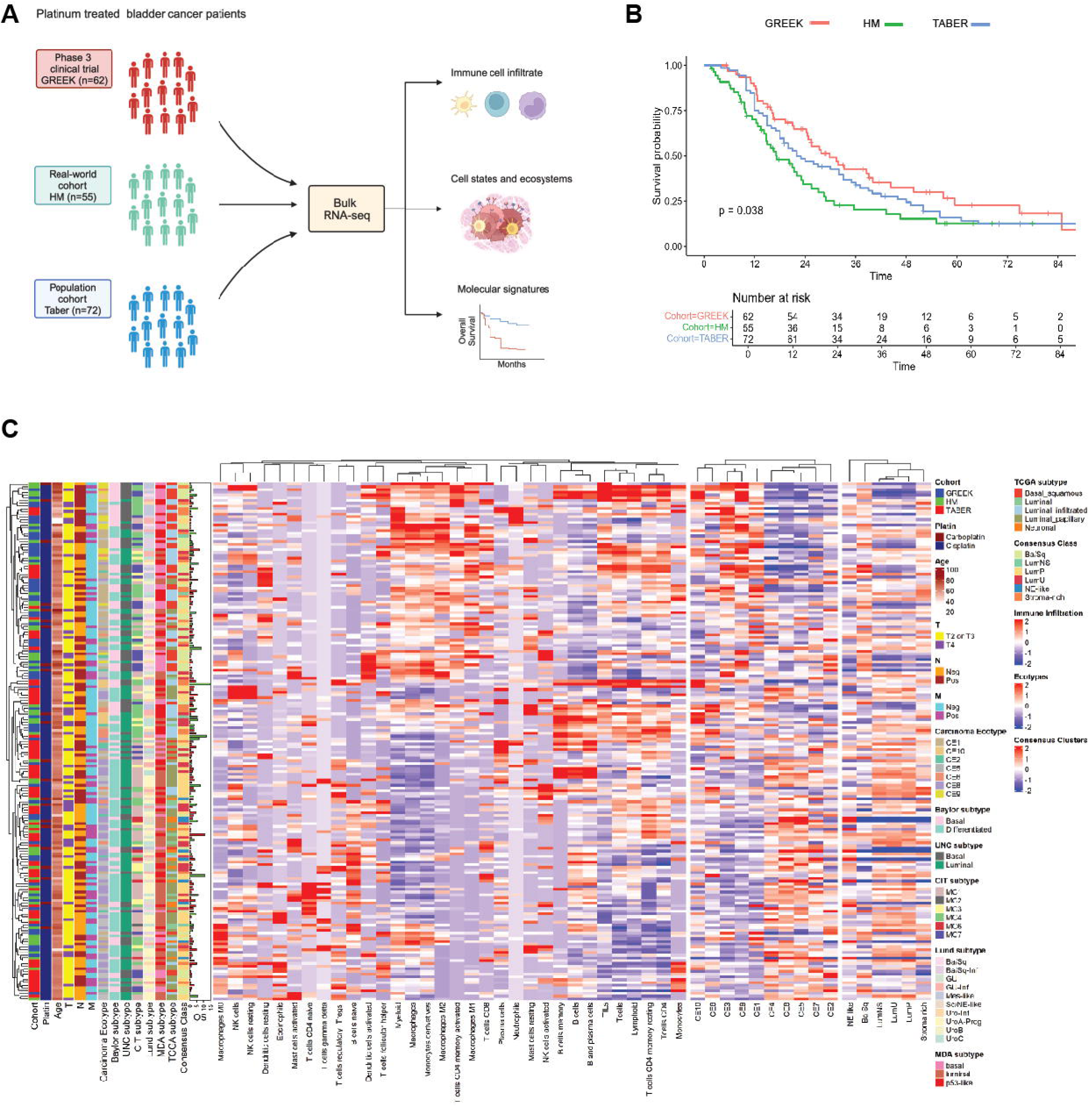
**Study design, methodology and cohort characteristics.** (A) Overview of the study workflow. (B) Kaplan-Meier analysis of overall survival (OS) in three independent cohorts used in the study: HM, GREEK, and Taber cohorts. (C) Heatmap displaying molecular tumor microenvironment (TME) characteristics across all samples. The heatmap includes assessments of tumor-infiltrating immune cells estimated using CIBERSORTx, scores for different cancer ecotypes (CEs) derived from EcoTyper, and scores for consensus molecular subtypes. Additional tracks display clinical characteristics and subtypes for each patient. Overall survival (OS) is provided in months; red bars indicate events (patients who died during follow-up), and green bars indicate censored observations (patients alive at last follow-up).

Due to the distinct inclusion criteria across the analyzed cohorts, we observed significant differences in overall survival across the cohorts (log-rank test, p = 0.038; Figure 1B, Supplementary Figure 1A). The HM cohort demonstrated significantly shorter OS than the GREEK cohort (HR 1.74; 95% CI: 1.13–2.68), but not the Taber cohort (HR 1.27; 95% CI: 0.86–1.90. Patients with metastatic disease at diagnosis also had significantly longer OS compared with patients diagnosed with locally-advanced disease (Supplementary Figure 1A–C). Moreover, patients treated with any regimen containing cisplatin had a trend toward improved survival compared to carboplatin-treated patients (HR 0.6; 95% CI: 0.40–1.12; p = 0.15; Supplementary Figure 1D). Treatment with MVAC did not confer a significant survival advantage over other cisplatin-based regimens (log-rank test, p = 0.059; Supplementary Figure 1E). Lastly, there was no association between survival time and age (Supplementary Figure 1F). Altogether, the five-year OS was poor across all cohorts, ranging from 13% to 23% (Table 1), consistent with reported outcomes for advanced urothelial cancer patients during the era of these studies.

### Lymphocyte infiltration was significantly associated with prolonged overall survival

We performed bulk RNA-sequencing (RNA-seq) for the GREEK and HM cohorts (Methods) and obtained publicly available bulk RNA-seq data for the Taber cohort. Following RNA-seq data harmonization, clinical and transcriptomic results were projected in a heatmap (Figure 1C), which included immune cell infiltration estimated with CIBERSORTx, tumor microenvironment (TME) ecotypes, and bladder cancer molecular subtypes based on the consensus classification (Methods).^14^ Unsupervised clustering of transcriptomic data did not segregate samples based on cohort, molecular subtype, TNM stage, or platinum-based regimen. However, several significant correlations were identified between the immune cell infiltration levels (Supplementary Figure 2). Immune cell subpopulation analysis revealed that high levels of lymphocytes in the primary tumor was significantly associated with prolonged OS after platinum-based chemotherapy (HR 0.34; 95% CI: 0.16–0.72; p = 0.005, Figure 2A). Notably, this effect was primarily driven by the presence of B cells rather than T cells (HR 0.14; 95% CI: 0.04–0.53; p = 0.004, Supplementary Table 2; Supplementary Figure 3). Among B cell subpopulations (which included naïve B cells and memory B cells), memory B cells were significantly associated with better OS (HR 0.20; 95% CI: 0.05–0.73; p = 0.015). Importantly, the estimated levels of these cell subgroups did not significantly differ among cohorts (Figure 2B). In contrast, the presence of myeloid cells, specifically macrophages, resting and activated dendritic cells, and neutrophils were associated with poor OS in platinum-treated urothelial cancer patients (Supplementary Figure 4).

**Figure 2.**
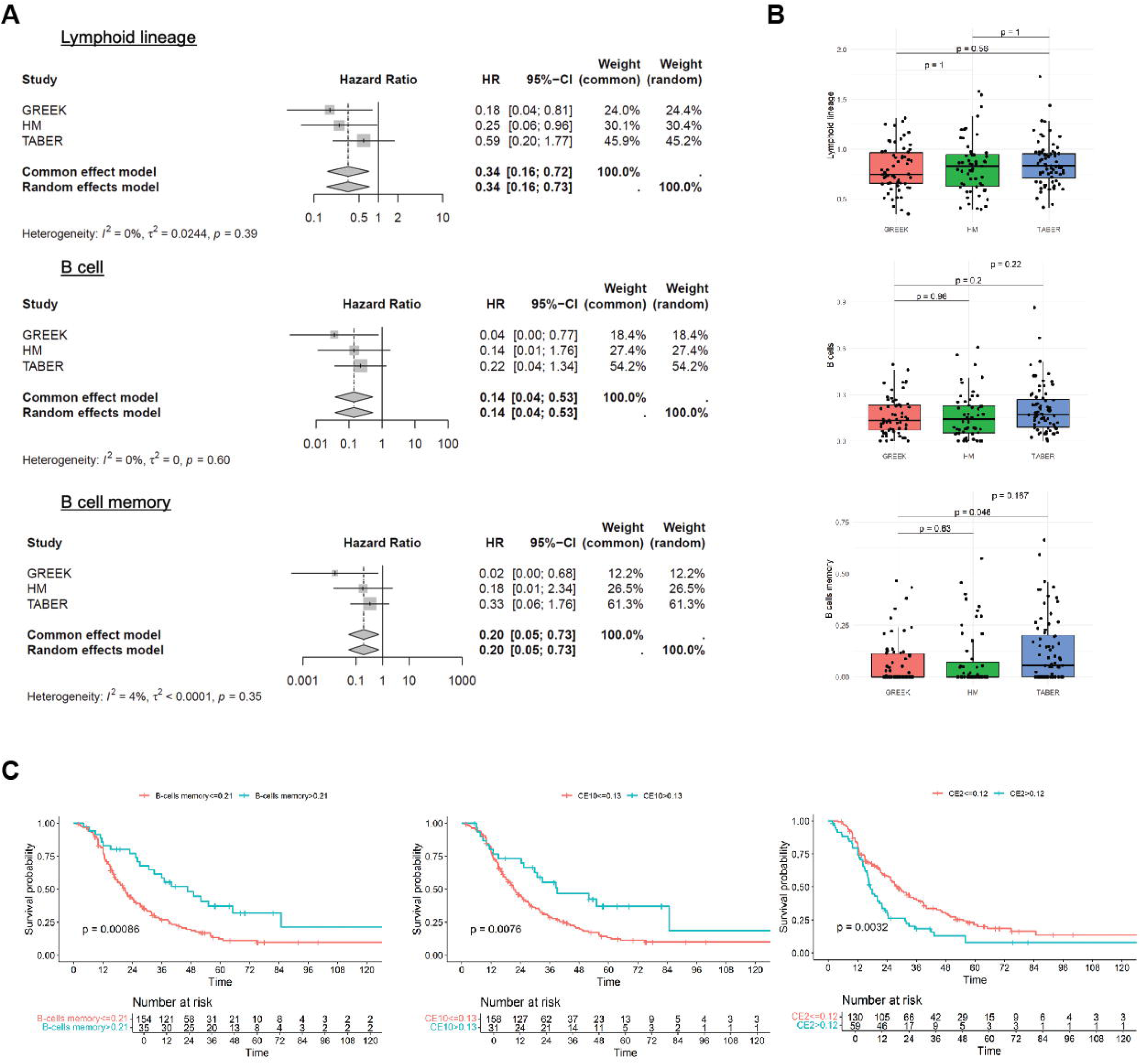
**Meta-analysis of tumor-infiltrating immune cells.** (A) Forest plots with hazard ratios (HRs) for overall survival, adjusted for age and T and M stages from the TNM classification. HRs for immune cell populations are estimated using CIBERSORTx. A fixed-effects model was used when the heterogeneity statistic (I²) was less than 50%. (B) Boxplots showing the distribution of lymphoid lineage cells, B cells, and memory B cells across the three cohorts. No significant differences were observed in the levels of these cell populations among the cohorts, indicating consistency across studies (Wilcoxon rank sum test). (C) Kaplan-Meier survival analysis for patient subgroups defined by high and low levels of memory B-cell infiltration, CE10 and CE2 ecotype score. Subgroups were determined using the maximally selected rank statistics (“maxstat”) method (Methods section). All comparisons showed statistically significant differences in overall survival, with higher scores associated with improved OS.

To further dissect how multicellular communities are correlated with OS, we applied EcoTyper^14^ across tumor samples from the three cohorts. Ecotypes (CE) CE2, CE6 and CE10 were significantly associated with prolonged OS, with an HR of 0.01 for CE10 (95% CI: 0.00–0.80; p = 0.040; Supplementary Figure 5 and 6). CE10 is a proinflammatory ecosystem with high B cell infiltration in the tumor microenvironment (Supplementary Figure 7&8). The ecosystem CE6 is highly specific for neoplastic tissues, which is consistent with the tumor samples profiled in this study, and CE2 is characterized by the presence of basal-like epithelial cells associated with resistance to cisplatin chemotherapy^9^.

Using cases from the 3 cohorts, we developed cutoffs to define of tumors with high memory B cell infiltration (Methods). Memory B cell-rich tumors comprised approximately 20% of all tumors, were evenly distributed across cohorts (chi-squared test; p = 0.177) and were associated with significantly better prognosis (Figure 2C; HR 0.47; 95% CI: 0.29–0.74). Moreover, high memory B cell presence was associated with CE10 (chi-squared test; p < 0.001) and CE2 (chi-squared test; p = 0.003) ecosystems consistent with EcoTyper expected immune cell infiltration.

Together, these results indicate the presence of memory B cells and a B cell-rich ecosystem in primary bladder tumors is associated with improved OS in patients with advanced bladder cancer treated with systemic platinum-based chemotherapy.

### Proinflammatory tumor microenvironment is associated with prolonged overall survival following cisplatin treatment

To investigate if cisplatin and carboplatin differentially modulate the bladder tumor microenvironment and how these changes impact OS, we divided the cohorts based on the treatment regimen (Supplementary Table 3). From the 189 patients included in the study, 167 patients received cisplatin-based and 22 received carboplatin-based chemotherapy. Increased tumor-infiltrating lymphocytes (TILs) were associated with significantly better OS in patients treated with cisplatin, but not in those receiving carboplatin (subgroup difference p = 0.019; Figure 3A). We further investigated which cell types are most strongly associated with better OS in cisplatin-treated patients and this effect was primarily driven by lymphoid lineage cells (subgroup difference p = 0.043), particularly memory B cells (subgroup difference p = 0.027, Supplementary Figure 9). We then defined differences in ecosystems in patients treated with cisplatin versus carboplatin and found CE10 (proinflammatory and B cell-rich) ecosystem had significant subgroup differences favoring cisplatin treated patients (Common effect model, HR 0.01; 95% CI: 0.00–1.62, Supplementary Figure 10A). Additionally, intergroup differences were noted for neutrophils and the LumNS consensus subtype in line with the fact that LumNS tumors displayed elevated stromal infiltration signatures, primarily fibroblastic, compared with other luminal tumors (Supplementary Figure 10B). Overall, these results indicate that immune cell infiltration, particularly of memory B cells and the proinflammatory CE10 ecotype, are associated with prolonged OS following cisplatin but not carboplatin chemotherapy treatment.

**Figure 3.**
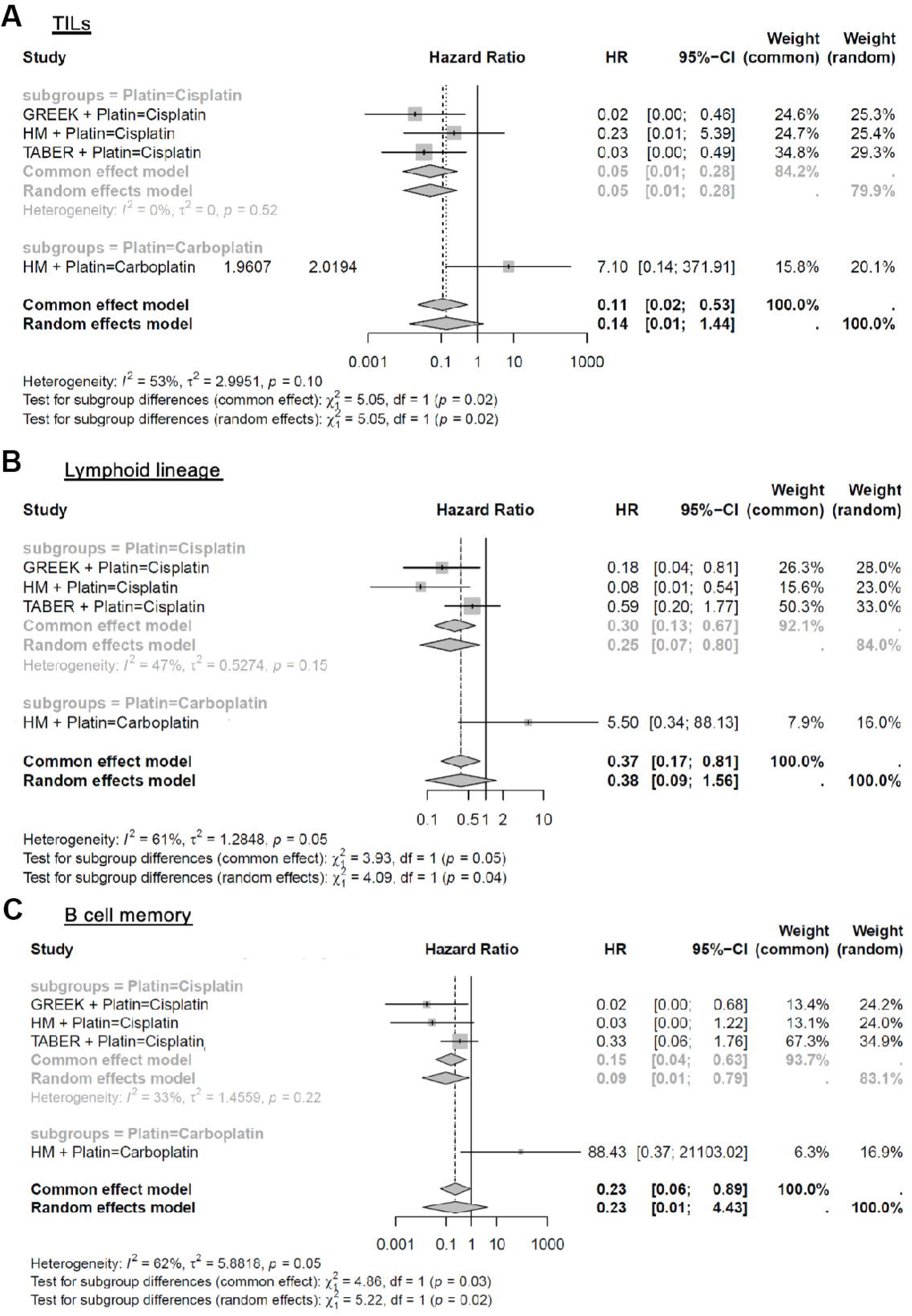
**Comparison of cisplatin versus carboplatin treatment on the association between immune cell infiltration and overall survival.** Forest plots display hazard ratios (HRs) for overall survival associated with tumor-infiltrating lymphocytes (TILs) (A), lymphoid lineage cells (B), and memory B cells (C), adjusted for age and T and M stage. A fixed-effects model was used when the heterogeneity statistic (I²) was less than 50%. Significant differences in HRs between the cisplatin and carboplatin groups were observed, confirmed by tests for subgroup differences (interaction p-values), indicating that immune cell infiltration was prognostic in cisplatin-treated patients but not in those treated with carboplatin.

### B-cell gene signatures serve as prognostic markers of survival

To build upon our observation that memory B cells were associated with improved OS, we investigated the association of other B cell-related gene expression signatures with predictive and prognostic implications across several cancer types. By calculating the scores using models developed in the original publications, we found that a B cell-related gene signature consisting of nine cytokine signaling genes^15^ was strongly correlated with memory B-cell infiltration in the tumor (Figure 4A; chi-squared test; p = 0.007). A B cell lineage depletion signature associated with poor survival in lung adenocarcinoma^16^ also showed worse OS in our bladder cancer cohort (Figure 4B; chi-squared test; p < 0.001). Moreover, another prognostic signature related to B-cell infiltration in pancreatic adenocarcinoma^18^ also showed a significant correlation with memory B-cell estimates (Figure 4C). Similar observations were noted for a 14-gene B-cell immune signature in triple-negative breast cancer derived from a pooled analysis of seven studies^17^. Here, the 14-gene signatures were also not significantly associated with B-cell memory (p = 0.083), but allowed for significant risk stratification, with a higher score associated with improved OS (Figure 4D).

**Figure 4.**
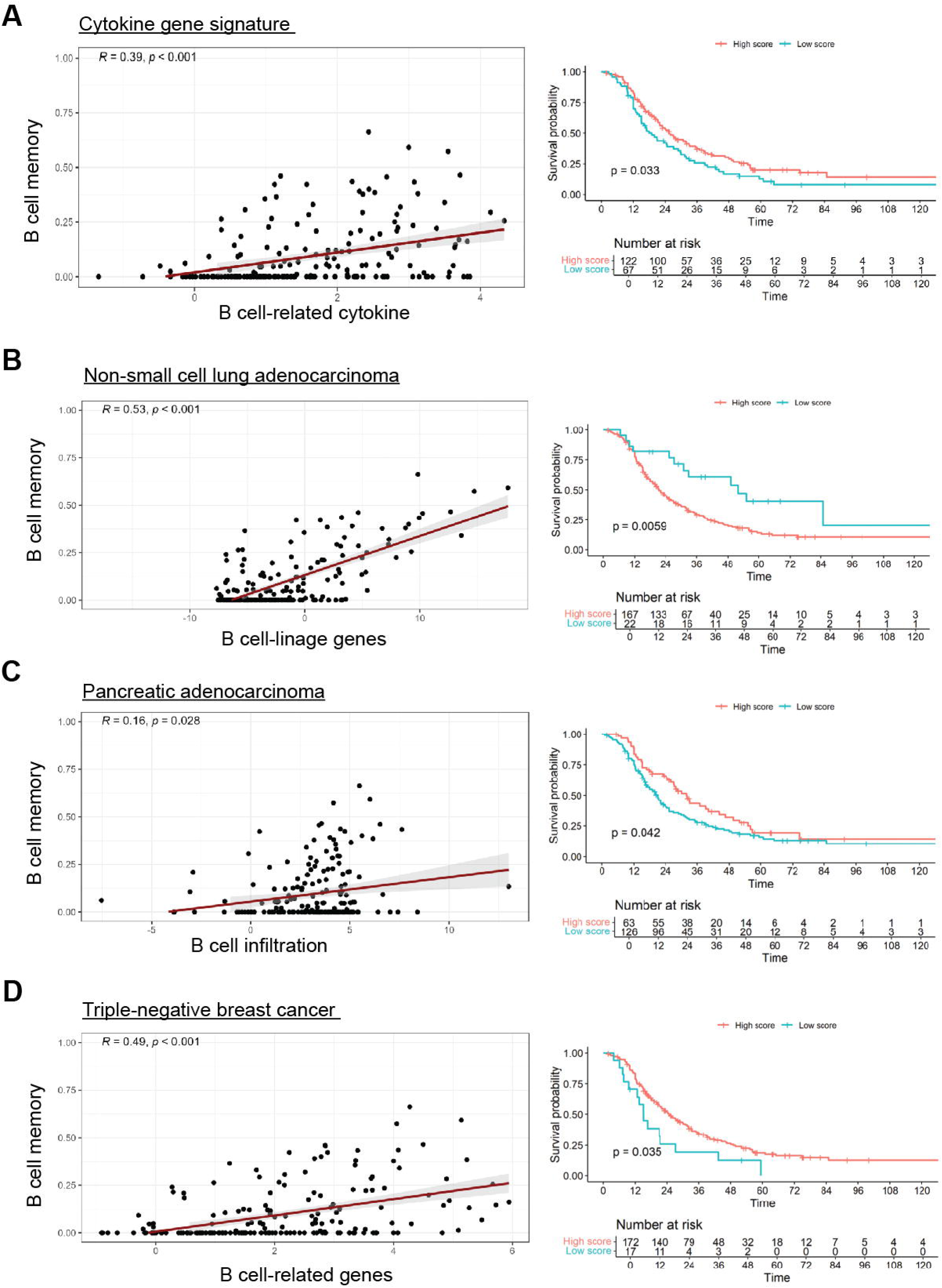
Concordance of select B-cell-related gene expression signatures with memory B-cell infiltration estimates and their prognostic abilities. Each panel represents one of four selected B-cell-related gene expression signatures (Methods section). All signatures showed significant correlation with memory B-cell infiltration estimates, affirming their relevance in our cohorts. (A) A cytokine gene signature demonstrated a positive correlation with memory B-cell infiltration and allowed significant stratification of overall survival, with higher scores associated with prolonged OS. (B) The B lineage-associated risk gene signature showed an inverse correlation with memory B-cell infiltration. Higher scores were associated with worse outcomes. This inverse relationship is consistent with the original publication where higher scores were associated with worse outcomes in non-small cell lung adenocarcinoma patients. (C) A prognostic gene signature related to B-cell infiltration in pancreatic adenocarcinoma correlated positively with memory B-cell estimates and significantly improved overall survival. (D) The B-cell immune signature derived from triple-negative breast cancer where higher scores were associated with better survival.

Overall, these results indicate that B cell gene expression signatures used to risk stratify patients in other cancer types also have prognostic implications in our bladder cancer cohorts. Additionally, three out of four signatures^15–17^ confirmed the significant differences in prognosis between cisplatin- and carboplatin-treated patients (Supplementary Figure 11). Pretreatment B-cell-related genes in the tumor were associated with worse prognosis in patients treated with carboplatin and better OS in cisplatin treated patients, suggesting a specific relationship between B cells and the effects of cisplatin in the immune microenvironment.

## Discussion

In this study, we investigated the prognostic significance of immune cell infiltration and molecular signatures in patients with locally advanced, surgically incurable, or metastatic urothelial bladder cancer treated with platinum-based chemotherapy. By analyzing pretreatment tumor samples from three independent cohorts, we found that infiltration of lymphocytes, particularly memory B cells, was associated with prolonged overall survival in patients receiving cisplatin-based chemotherapy. In contrast, this association was not observed in patients treated with carboplatin-based chemotherapy.

Our findings suggest that the presence of memory B cells in the bladder cancer tumor microenvironment may enhance the efficacy of cisplatin-based chemotherapy. Recent studies have increasingly recognized the significant role of B cells in anti-tumor immune responses, particularly their involvement in tertiary lymphoid structures^19^. B cells and plasma cells contribute to tumor suppression through classical humoral immunity mechanisms, for example antibody-dependent cell cytotoxicity and complement activation, as well as presenting antigens and producing cytokines that enhance T cell responses. Tertiary lymphoid structures provide a microenvironment where B cells can effectively present antigens, leading to optimal activation of T cells and the generation of effector and memory B cells^20^. Recent studies reinforce tumor-infiltrating B cell-mediated antitumor immunity through antigen presentation to T cells; their role in assembling and perpetuating immunologically ‘hot’ tumor microenvironments involving T cells and natural killer cells; and their potential to immune editing and be effective against tumors with heterogenous cell types^21^.

Our results also suggest that the presence of memory B cells in the bladder cancer tumor microenvironment may enhance the efficacy of cisplatin chemotherapy but not carboplatin. Cisplatin has been shown to induce immunogenic cell death, leading to the release of tumor antigens and the activation of dendritic cells and T cells^22^. Our findings support this antitumor cell death mechanism, as markers of T cell activation (e.g., CD69, IFNG, CD25/IL2RA) and dendritic cell activation (e.g., CD40, CD80, CD86, CCR7) were significantly upregulated in specimens with high memory B-cell infiltration (Supplementary Figure 12). This suggests that cisplatin’s therapeutic benefits may be partly mediated through its immunomodulatory effects, which are amplified in tumors with higher lymphocyte infiltration. Conversely, carboplatin may not elicit the same immune responses, potentially explaining the lack of association between immune infiltration and survival in carboplatin-treated patients. A recent subanalysis of IMvigor130^5^ showed that combination of gemcitabine and cisplatin versus gemcitabine and carboplatin with atezolizumab induces transcriptional changes in circulating immune cells, including upregulation of antigen presentation and T cell activation programs. In the same study, *in vitro* experiments demonstrated that cisplatin, but not carboplatin, exerted direct immunomodulatory effects on cancer cells, promoting dendritic cell activation and antigen-specific T cell killing. Our results also support that cisplatin modulates the microenvironment in the primary tumor with a positive impact in prolonged overall survival.

We provided further insights in the complex cellular composition of the tumor microenvironment of primary pretreatment bladder cancer and the implications in overall survival. The identification of specific ecosystems, such as CE10—which is characterized by proinflammatory and lymphocy-rich features—provides additional evidence into the prognostic significance of immune cell populations. The correlation between memory B-cell presence and the CE10 ecotype further emphasizes the role of adaptive immunity in influencing patient outcomes. Ecotype CE10 is becoming one of the most well established predictive TME factors associated with immunotherapy across cancer types^23^. Our results are in line with these observations where high number of B cells are associated with favorable prognosis and improved overall survival in patients treated with cisplatin-based chemotherapy. Thus, our study implicates a role for memory B cells in bladder cancer as good indicators of overall survival that can be harnessed as predictive biomarkers of response to cisplatin-based chemotherapy and open the possibility of B cell-directed therapies in combination with chemo-immune checkpoint inhibitors in future clinical trials.

Interestingly, we were not able to notice noteworthy associations between prognosis, immune cell infiltration and consensus molecular clustering. Mixed validation results still hinder translation of molecular subtypes to clinic, with new classification systems being constantly developed and improved and ongoing clinical trials^24^. Finally, the validation of our findings using B-cell-related gene expression signatures from other cancers strengthens the evidence for the prognostic value of B-cell infiltration. The consistency across different signatures and cohorts suggests that these gene expression profiles could serve as robust biomarkers for patient stratification and treatment planning.

This study has a few limitations. First, the retrospective design and the use of archival samples introduce the potential for selection bias and limit the ability to establish true causal relationships. Although we adjusted for confounding variables such as age, tumor stage and metastatic disease, unmeasured factors like performance status, metastatic location, tumor burden and comorbidities could influence survival outcomes and were not fully accounted for. Second, the heterogeneity among the three cohorts—differences in patient characteristics, treatment regimens, and duration of follow-up— may affect the generalizability of our findings. The small sample size in the carboplatin-treated subgroup limits the statistical power to detect significant differences and may influence the robustness of subgroup analyses. Third, the reliance on computational estimations of immune cell populations from bulk RNA-seq data, while informative, may not capture the full diversity of the tumor microenvironment. Techniques like single-cell RNA-seq could provide more precise measurements of immune cell infiltration, but these are very costly approaches that are difficult to implement in large cohorts as the ones profiled in our study. Finally, potential batch effects that characterize RNA sequencing studies might occur, but we used standardized pipelines and meta-analysis approaches to mitigate residual inconsistencies that might impact the results.

In conclusion, our study highlights a significant association between memory B-cell infiltration and prolonged OS in patients with urothelial bladder cancer treated with cisplatin-based chemotherapy. These findings suggest that immune cell infiltration, particularly by memory B cells, could serve as a prognostic biomarker to guide treatment selection and optimize therapeutic outcomes. The differential effects observed between cisplatin and carboplatin underscore the importance of considering the tumor immune microenvironment when selecting chemotherapy regimens. Understanding the interplay between cytotoxic therapies and the immune microenvironment can biologically inform future treatment approaches in urothelial cancer.

## Methods

### Patient cohorts

We analyzed pretreatment bladder tumor tissue (from bladder) from a phase III clinical trial (GREEK cohort, ACTRN 12610000845033), a real-word bladder cancer patient cohort (HM cohort) and previously published results from a cohort of 300 bladder cancer patients ^9^ referred to as the TABER cohort, from which only patients with locally advanced and metastatic cancer (surgically uncurable) were included in the analysis. Our study included patients with locally advanced, surgically unresectable, or metastatic urothelial bladder cancer who were treated with either cisplatin- or carboplatin-based chemotherapy. Patients who did not receive platinum-based chemotherapy or who received neoadjuvant chemotherapy before surgery were excluded from the analysis.

Clinical data, including diagnostic staging, treatment details, and follow-up information, were collected for all cohorts. Biopsies from the HM and GREEK cohorts were processed for bulk RNA sequencing (RNA-seq) and publicly available RNA-seq data for the Taber cohort was included in the analysis.

The GREEK cohort includes patients from the HR 16/03 trial (ACTRN12610000845033) —a prospective, open-label, randomized phase III study comparing two dose-dense chemotherapy regimens: methotrexate, vinblastine, doxorubicin, and cisplatin (DD-MVAC) versus gemcitabine and cisplatin (DD-GC) in patients with inoperable, metastatic, or relapsed urothelial cancer ^10^ Patients with good performance status (ECOG 0-1) were randomly assigned to receive either DD-MVAC or DD-GC. The median follow-up was 52.1 months (89 events). The study concluded that DD-GC was not superior to DD-MVAC but was better tolerated.

The HM cohort comprised a prospectively observed group of real-world patients with locally advanced, surgically incurable, or metastatic urothelial bladder cancer diagnosed and treated at Hospital del Mar (Barcelona, Spain) from 2005 to 2010, with follow-up until 2012. Patients in this cohort were primarily treated with platinum-based chemotherapy.

From the Taber cohort, we included only patients who received first-line chemotherapy upon detection of locally advanced (T4b) or metastatic disease. In the original publication, cisplatin-based chemotherapy was administered in approximately 98% of patients.

In all cohorts, pretreatment staging was based on baseline imaging and the pathological assessment on the transurethral resection of bladder tumor (TURBT) or radical cystectomy specimens. Treatment response was defined as complete response (CR) or partial response (PR) based on post-treatment cross-sectional imaging according to the RECIST 1.1 guidelines. Overall survival (OS) was defined as the time from muscle-invasive bladder cancer (MIBC) diagnosis to death or end of follow-up. The clinical data used in this analysis are summarized in Supplementary Table 1.

### Bulk RNA-sequencing

Total RNA was extracted from formalin-fixed, paraffin-embedded (FFPE) bladder tumor samples using the RNeasy Mini Kit (Qiagen) according to the manufacturer’s instructions. RNA quality and integrity were assessed using an Agilent 2100 Bioanalyzer, ensuring RNA integrity numbers (RIN) greater than 7.0. Sequencing libraries were prepared using the Illumina TruSeq Stranded mRNA Library Prep Kit. Libraries were sequenced on an Illumina NextSeq 500 platform to generate paired-end reads of 75 base pairs in length.

Raw sequencing data were processed using the nf-core/rnaseq pipeline (version 3.14.0; https://nf-co.re/rnaseq/3.14.0/), implemented with Nextflow for reproducibility and scalability ^11^. Briefly, initial quality control of raw reads was performed using FastQC. Adapter sequences and low-quality bases were trimmed using Trim Galore!, which incorporates Cutadapt. Cleaned reads were aligned to the human reference genome (GRCh38) using the STAR aligner. Gene expression quantification was conducted using featureCounts, providing gene-level counts for downstream differential expression analysis. Additionally, transcript-level quantification was performed using Salmon in quasi-mapping mode, enabling fast and accurate estimation of transcript abundances directly from the trimmed reads. Differential gene expression analysis was carried out using DESeq2 within the R statistical environment.

### Immune cell infiltrates and cellular communities

An overview of the analyses performed is presented in Figure 1A. To investigate the impact of the immune tumor microenvironment (TME) on overall survival in patients treated with platinum-based chemotherapy, we employed CIBERSORTx ^12^ in absolute mode to deconvolute the abundance of immune cell populations in each pretreatment tumor sample. CIBERSORTx utilizes gene expression signatures to characterize cellular heterogeneity from bulk tissue transcriptomic data without the need for physical cell isolation. In absolute mode, cellular fractions are scaled to scores that reflect each cell type’s absolute proportion, allowing for comparison across both samples and cell types. As described in previous studies, the tumor-infiltrating lymphocyte (TIL) score for each sample was determined by summing the estimated proportions of all immune cell types except eosinophils and neutrophils ^13^. Monocyte derivatives were analyzed by adding estimates for macrophages (M0, M1, and M2), dendritic cells (both activated and resting dendritic cells), and monocytes.

We then used EcoTyper ^14^ to systematic identify cell states and cellular communities in tumor samples. Cell types are first deconvolved from bulk RNA-seq data to estimate their proportions within each sample and cell states defined based on gene expression profiles using non-negative matrix factorization (NMF). By analyzing the co-occurrence patterns of these cell states across multiple samples, EcoTyper clusters cell states into recurrent multicellular communities termed “cellular ecotypes” (CEs). Each CE represents a specific configuration of cell states that reflects the functional organization of the tissue microenvironment.

Briefly, EcoTyper ^14^ characterizes 10 ecotypes. CE1-high tumors are enriched in fibroblasts, depleted in lymphocytes and strongly associated to a higher risk of cancer-associated death. CE2-high tumors are distinguished by elevated levels of basal-like epithelial cells, are also lymphocyte-deficient and associated with a significantly increased risk of mortality. CE6-high tumors are unique because there are no immune cells as opposed to CE9-high tumors where proinflammatory immune cells, interferon-gamma (IFN-γ) activation that indicates a robust immune response association with prolonged OS. Lastly, CE10-high tumors also have high levels of proinflammatory cells, are strongly associated with longer overall survival, but are distinguished by a higher B-cell content.

After defining the infiltrating immune cell populations, we used the consensus molecular classification of muscle-invasive bladder cancer^25^ to integrate tumor-intrinsic features with the surrounding microenvironment and comprehensively characterized the bladder cancer ecosystem. Consensus clustering was implemented using the “BLCAsubtyping” R package.

### Statistical analysis

Intragroup associations were assessed using standard statistical tests appropriate for the variable types. For comparisons of categorical variables, the chi-squared test was used. Survival analyses incorporated the log-rank test and Cox proportional hazards regression.

Due to potential batch effects and differences in sequencing depth across cohorts, we decided to analyze CIBERSORTx results using a meta-analysis approach. For each cohort, we performed multivariable Cox proportional hazards regression analyses to evaluate the association between immune cell infiltration and overall survival. The Cox models were adjusted for potential confounders, including age at diagnosis, T stage, and M stage. The primary variable of interest in each model was the estimated infiltration level of a specific immune cell type. Hazard ratios (HRs) with 95% confidence intervals (CIs) and p-values were calculated for each immune parameter.

To synthesize results from individual cohorts, we conducted meta-analyses for each immune parameter. Log-transformed HRs and their standard errors were calculated from the Cox regression outputs of each cohort. Both fixed-effect and random-effects meta-analyses were performed to obtain pooled HRs. Heterogeneity among studies was assessed using Cochran’s Q test and quantified with the I² statistic. We reported pooled HRs with 95% CIs, p-values for overall effects, and heterogeneity statistics. Forest plots were generated to illustrate the HRs from individual cohorts and the pooled estimates.

To investigate the potential differential impact of cisplatin versus carboplatin on the association between immune cell infiltration and overall survival, we performed subgroup analyses. Patients were stratified based on the platinum agent received (cisplatin or carboplatin). Multivariable Cox proportional hazards models were repeated within each platinum agent subgroup for each immune parameter. Meta-regression analyses were conducted to assess whether the type of platinum agent modified the effect of immune cell infiltration on survival outcomes. Subgroup-specific HRs, pooled HRs within subgroups, and interaction p-values from meta-regression analyses were reported.

We evaluated potential publication bias using funnel plots and statistical tests, including Begg and Mazumdar’s rank correlation test and Egger’s regression test. As all comparisons were planned and limited to chosen comparisons, no p-value adjustment was applied at this step.

To visualize the prognostic abilities of selected parameters, we employed the maximally selected rank statistics method for each parameter to determine the cut point that most significantly separated patients into groups with different overall survival outcomes. This non-parametric approach identifies the value of a continuous variable that maximizes the difference in survival between two groups. After determining the optimal cut points, patients were divided into high and low groups for each immune parameter. Kaplan-Meier survival curves were generated to compare overall survival between the high and low groups, and the log-rank test was used to assess the statistical significance of differences between survival curves. Univariate Cox regression analyses were conducted to estimate hazard ratios and 95% confidence intervals for the association between group membership (high vs. low) and overall survival.

Additionally, we investigated associations between the immune parameter groups and various clinical and molecular variables. Associations with categorical variables were evaluated using chi-squared tests, while continuous variables were compared using t-tests or non-parametric equivalents, depending on data distribution.

### Gene expression signature analysis

To validate the potential prognostic implications of B cells in bladder cancer, we assigned four B-cell function gene expression signatures for each tumor: (i) cytokine signaling pathways pertinent to B-cell function^15^ (ii) B-cell lineage gene signature^16^ (iii) lymphocyte maturation, activation, differentiation ^17^ (iv) signature associated with B-Cell infiltration in human cancer^18^. For each signature, we calculated a score for every patient according to the methodologies described in the original publications. For signatures defined by the mean expression of included genes, we calculated the average transcripts per million (TPM) value of those genes for each patient. For signatures with specified coefficients, we computed a weighted sum by multiplying each gene’s expression level by its corresponding coefficient from the original publication.

Due to the inherent distribution of the data, Spearman’s rank correlation coefficients were calculated between each signature score and the estimated memory B-cell levels to determine the strength and direction of the associations. We employed the maximally selected rank statistics method to identify optimal cutpoints for stratifying patients based on their gene signature scores, identifying thresholds that most significantly differentiated patient survival outcomes without prior specification of the cutoff point. Subsequent analyses followed the same methodology as for the immune-related parameters, including survival analyses and assessments of associations with clinical and molecular variables.

## Data and code availability

Code and raw data files will be made publicly available at the time of publication.

## Acknowledgments

We thank the patients who participated in this study. This work was supported by NCI K08CA282969-01A1 (to F.L.F.C.), Bladder Cancer Advocacy Network Young Investigator Award and Career Development Award (to F.L.F.C.), NCI 3P30CA006516-59W2 Early Stage Surgeon-Scientist Program award (to F.L.F.C.), NCI R01CA279221 (to K.W.M).

## Declaration of interests

J.B. reports the following financial interests: advisory board participation with AstraZeneca, BMS, Merck, and Pfizer; participation as an invited speaker or lecturer by Merck and MSD; stocks and/or shares from Bicycle; and royalties from UpToDate.

**Table 1. Baseline clinical characteristics and survival outcomes of the three study cohorts.** The table summarizes patient demographics, TNM staging at diagnosis, treatment details, and overall survival metrics for the GREEK, HM, and TABER cohorts. The GREEK and TABER cohorts exclusively received cisplatin-based chemotherapy, whereas 40% of HM patients were treated with carboplatin-based regimens. Significant differences were observed in age at diagnosis (p = 0.032) and metastatic disease at diagnosis (M+ vs. M0, p = 0.032). Median survival was highest in the HM cohort (29.8 months) and lowest in the GREEK cohort (17.2 months). Five-year survival rates were poor across all cohorts, ranging from 13% (HM) to 23% (GREEK).

## Supplementary information

**Supplementary Table 1.** Clinical characteristics of included patients. TNM at the time of diagnoses.

**Supplementary Table 2.** Meta-analysis results for all established parameters.

**Supplementary Table 3.** Meta-analysis results for all established parameters with assessment of difference between cisplatin and carboplatin-treated patients.

**Supplementary** Figure 1. Panels A-C show univariable overall survival analysis of TNM characteristics. P-value provided on the plot was calculated using log-rank test. Metrics presented in the table derived from Cox regression. For age of diagnosis the cutoff for grouping was determined using maxstat R package to maximize log-rank statistic with p-value approximation based on an improved Bonferroni inequality (pLausen94).

**Supplementary** Figure 2. Network plot of significant correlations between molecular metrics. This network visualizes the significant correlations between parameters (correlation coefficient |r| > 0.4, p < 0.05). Nodes represent individual immune cell populations or molecular metrics, while edges represent significant positive (red) or negative (blue) correlations between them. The thickness of the edges corresponds to the strength of the correlation, with thicker lines indicating stronger associations.

**Supplementary** Figure 3. Forest plot of meta-analysis evaluating the association between T-cell infiltration and overall survival. The figure shows hazard ratios (HRs) and 95% confidence intervals (CIs) for T-cell infiltration in each cohort and the pooled estimate, indicating no significant association with survival.

**Supplementary** Figure 4. Forest plots of meta-analysis evaluating the association between myeloid cell infiltration and overall survival. The figure displays hazard ratios (HRs) and 95% confidence intervals (CIs) for monocytes derivatives and neutrophils in each cohort and the pooled estimates, indicating an association with poorer survival.

**Supplementary** Figure 5. Kaplan-Meier analysis of overall survival for different Ecotypes. Log-rank test showed borderline significance for differences between ecotypes.

**Supplementary** Figure 6. Association between tumor ecotypes and overall survival. This figure presents hazard ratios (HRs) and 95% confidence intervals (CIs) for Ecotypes CE2, CE6, and CE10 derived from EcoTyper analysis, indicating their significant associations with overall survival.

**Supplementary** Figure 7. Correlations between B cell memory infiltration and CE10, CE7 and CE6 scores.

**Supplementary** Figure 8. Comparison of immune cell infiltration levels in tumors with high versus low memory B-cell infiltration. The figure shows boxplots of estimated immune cell proportions, demonstrating increased lymphoid infiltration, particularly CD4⁺ T cells, in tumors with high memory B-cell infiltration.

**Supplementary** Figure 9. Higher B-cell infiltration was not significantly associated with worse OS in patients receiving carboplatin.

**Supplementary** Figure 10. Differences in immune cell infiltration and molecular subtypes scores between cisplatin- and carboplatin-treated patients. This figure illustrates the differences in Ecotype CE10 scores (A) and Luminal Non-Specified (LumNS) molecular subtype (B) score between the two treatment groups.

**Supplementary** Figure 11. Prognostic significance of B-cell related gene signatures in cisplatin-versus carboplatin-treated patients. This plot depicts the forest plots for each signature with results of the test for subgroup differences.

**Supplementary** Figure 12. Differential expression of immune activation markers in tumors with high versus low memory B-cell infiltration. Boxplots display the expression levels of T-cell activation markers (CD69, IFNG, IL2RA) and dendritic cell activation markers (CD40, CD80, CD86, CCR7) in tumors stratified by high and low memory B-cell infiltration. Significant upregulation of CD69 (p = 0.00017) and CCR7 (p = 0.0016) is observed in tumors with high B-cell memory infiltration. Trends toward increased expression of IL2RA (p = 0.053), CD40 (p = 0.16), and CD86 (p = 0.14) suggest enhanced immune activation, supporting the role of memory B cells in modulating the tumor microenvironment in response to cisplatin treatment.

